# GPCR-Nexus: Multi-Agent Orchestration for Knowledge Retrieval

**DOI:** 10.64898/2026.04.10.696782

**Authors:** Jackson Spieser, Juechen Yang, Jarek Meller, Krushna Patra, Behrouz Shamsaei

**Affiliations:** University of Cincinnati, College on Medicine Cincinnati, Ohio; Department of Biostatistics, Health Informatics and Data Sciences, University of Cincinnati College of Medicine, Cincinnati, Ohio; Division of Biostatistics and Bioinformatics, Department of Environmental and Public Health Sciences, University of Cincinnati College of Medicine, Cincinnati, Ohio; Division of Biomedical Informatics, Cincinnati Children’s Hospital Medical Center, Cincinnati, Ohio; Department of Cancer Biology, University of Cincinnati College of Medicine, Cincinnati, Ohio; Institute of Engineering and Technology, Faculty of Physics, Astronomy and Informatics, Nicolaus Copernicus University, Torun, Poland; Department of Computer Science, University of Cincinnati College of Engineering and Applied Sciences, Cincinnati, Ohio

## Abstract

We present GPCR-Nexus, an AI-driven platform for integrated exploration of G protein–coupled receptor (GPCR) biology that unifies structured databases with unstructured scientific literature. The system combines a GPCR–ligand knowledge graph with vector-based semantic retrieval to enable comprehensive, up-to-date information access. Central to GPCR-Nexus is a multi-agent architecture in which specialized components coordinate query planning, evidence retrieval, validation, and synthesis. This design ensures that generated responses are grounded in verifiable sources while maintaining coherence across heterogeneous data modalities. By jointly leveraging curated databases and primary literature, GPCR-Nexus enables context-aware reasoning over molecular interactions, functional mechanisms, and disease associations. The platform produces citation-backed outputs with traceable evidence, addressing limitations of conventional database queries and standalone language models. We detail the system architecture, data integration strategy, and agent orchestration framework, and demonstrate its utility through representative query scenarios. GPCR-Nexus provides a scalable approach to combining structured and unstructured biomedical knowledge using agent-based AI, offering improved accuracy, interpretability, and coverage. This work establishes a foundation for trustworthy, AI-assisted knowledge synthesis in GPCR research and drug discovery.

## Introduction and Motivation

GPCRs constitute the largest family of human cell-surface receptors and are major drug targets; roughly one-third of FDA-approved drugs act on GPCRs [1]. However, knowledge of GPCR-ligand interactions is highly fragmented. Databases like GPCRdb and ChEMBL curate sequences, structures, and bioactivity data, but they fail to provide the contextual narrative from literature [2,3]. The mechanistic roles of GPCRs (e.g. in disease) remain buried in unstructured articles, so researchers must cobble together information from multiple sources or rely on general LLMs (e.g. ChatGPT). However, studies show they can hallucinate answers or references, omit evidence, and have a training cutoff [16–19]. GPCR-Nexus (gpcr-nexus.org) was developed to fill this gap: it employs multi-AI agent orchestration for information retrieval and synthesizing a response that combines vector-based semantic search of the literature with a structured GPCR–ligand knowledge graph [4]. Multiple AI agents (source planner, fact reviewer, database, synthesizer) coordinate to ensure that answers are factual and citation-backed, delivering coherent, context-rich responses that span both databases and the latest publications. A comparison of standard LLMs, GPCR curated databases, and GPCR-Nexus is provided in Table 1. The contents of GPCR-Nexus are explained throughout the rest of this paper.

**Table 1:** Comparisons of GPCR-Nexus, Curated GPCR databases, and LLMs.

| Aspect | GPCR-Nexus | GPCRdb | ChEMBL | Standard LLMs |
| --- | --- | --- | --- | --- |
| **Focus / Scope** | GPCR-centric knowledge synthesis: literature-driven Q&A powered by AI agents | GPCR-specific curated repository: sequences, structures, ligands | Broad bioactivity repository: drug-like compounds across all targets, including GPCRs | General-purpose language understanding and generation; not domain-specialized |
| **Data Content** | Graph + text: GPCRs, ligands, pathways, diseases; literature-derived relations and snippets | Sequences + interactions: ~805 human GPCRs, 1,700 structures, 222k ligands | Compounds + bioactivity: >2.4 M compounds and 20 M measurements on 17k targets | Entire training corpus (books, web, code); no explicit structured bio or GPCR data |
| **Data Format** | Hybrid: structured knowledge graph + unstructured text (semantic embeddings) | Structured: predefined fields with curated integration | Structured: relational compound–target–activity schema | Unstructured text; knowledge implicit in model parameters |
| **Use of AI** | Extensive: multi-agent LLM orchestration (planning, retrieval, verification, synthesis) | None: manual curation and static web queries | None: curated only; users apply external AI tools | Extensive: single LLM for generation; no retrieval or verification pipeline |
| **Literature Integration** | Direct: full-text indexing and citation; NLP-extracted graph relations | Indirect: curated facts from literature; no live text integration | Indirect: curated activity records; no live text integration | Indirect: knowledge only via training data; cannot cite or retrieve actual passages |
| **Context & Hypothesis Support** | High: narrative answers, experiment suggestions, context-aware synthesis | Moderate: visual tools support interpretation; user-driven hypotheses | Low: data only; no contextual analysis or suggestions | Low: factual-only responses; prone to hallucinations; no experimental planning |
| **User Interface** | Natural language interface: multi-source answer with citations; interactive web portal with structured views, Sankey plots, diagrams; tabular interface with keyword/structure search | Static web UI: GPCR family browser, sequence alignments, ligand lists | Web UI: searchable tables of compounds and activities | Chat interface (text in/text out); no structured visualizations |
| **Query Flexibility** | Very high: understands multi-part natural language queries semantically | Limited: structured input only; no NL understanding | Moderate: advanced filters and chemical search, but no NL | Very high: natural language queries, but lacks semantic grounding |
| **Extensibility & Updates** | Automated and generalizable: ingests new literature continuously; adaptable to other domains | Manual but evolving: domain-focused expansions and user tools | Institutional-scale: regular updates by curators; high scalability | Fixed model checkpoint; retraining required for updates |
| **Strengths / Unique Features** | Contextual AI synthesis: literature + graph + agentic reasoning in one; domain-specific accuracy; curated visualizations; integrated hypothesis support | Domain-specific curation; reliable sequence and structure data | Breadth + depth of bioactivity data | Fluent, coherent text generation; broad general knowledge |
| **Limitations** | AI dependence: output quality tied to model; minimal graphics | No narrative: requires manual integration of context | No context: data only; user must interpret | Hallucinations; no citations; fixed knowledge cutoff; ungrounded answers |
| **Table 1: Comparisons of GPCR-Nexus, Curated GPCR databases, and LLMs** |  |  |  |  |

### Comparison of GPCR-Nexus to Existing Approaches

#### Comparison to Database-Centric Approaches

GPCRdb and ChEMBL provide high-quality curated data on GPCR sequences, structures, and ligand bioactivities, but only answer specific factual queries. They do not integrate findings across studies or explain the physiological context. For example, GPCRdb can list which ligands bind a receptor, but it cannot summarize *why* those ligands matter in a biological process. GPCR-Nexus overcomes this by retrieving relevant literature passages and linking them via a graph of entities and relations, so that a single query (“What does ligand Y do in cancer?”) yields a synthesized answer rather than isolated data points. Because these databases are purely structured, they omit the narrative insights found in papers. Much GPCR knowledge (mechanisms, pathways, disease relevance) resides in prose alongside the existing databases. GPCR-Nexus searches its indexed corpus of documents using embedding-based queries, an assembled knowledge graph, and a curated offline database. This distinct architecture provides responses that have mechanistic context along, the structured accuracy of a database, and the ability to return references. In this way, it effectively synthesizes information across sources, rather than requiring the user to search PubMed separately.

GPCRdb and ChEMBL must be manually curated and cannot easily incorporate the latest literature. In contrast, GPCR-Nexus continuously ingests new PDFs into Azure Blob storage, automatically chunking and embedding them. The chunks are indexed with OpenAI’s text-embedding-3-small model and stored using a Hierarchical Navigable Small World (HNSW) graph for fast approximate nearest-neighbor search [10]. This pipeline means new findings are quickly available for retrieval. Because GPCR-Nexus uses a hybrid vector and graph design, it yields higher recall and accuracy on complex queries than database-only or LLM-only approaches. Recent work on hybrid RAG confirms that combining knowledge graphs with vector retrieval improves answer quality [5,6]. In practice, GPCR-Nexus can answer integrative questions in seconds – a task that would otherwise take hours of manual searching through GPCRdb, ChEMBL, and the literature.

### Comparison to LLM-Based Tools

General-purpose LLMs such as ChatGPT, Claude’s Sonnet, and Google’s Gemini can produce remarkably fluent prose. Still, numerous studies demonstrate they lack grounding in an external knowledge base and frequently “hallucinate” information. In a systematic evaluation of scientific reference accuracy, ChatGPT-3.5 fabricated 39.6 % of its citations, GPT-4 hallucinated 28.6 %, and Bard (now Gemini) up to 91.4 % of the time when asked to replicate systematic review references [16].

Moreover, ChatGPT achieves only 58.5 % accuracy at distinguishing factual from hallucinated summaries, barely above chance [17]. Large-scale benchmarking (HaluEval) confirms that about 19.5 % of ChatGPT responses contain unverifiable content, and open-source LLMs perform similarly poorly without external knowledge support [18]. Critically, when ChatGPT is prompted to provide citations, only 14 % of its suggested references exist, and merely half of those support the claimed facts [19].

Search-oriented tools like Perplexity AI aim to address these flaws by surfacing source links; however, they still rely on third-party LLMs and suffer from accuracy, reliability, and recency issues [20]. Underlying most LLM-models is a fixed training cutoff, so they cannot incorporate the latest GPCR studies or other emerging biomedical research without manual retraining or the ability to access this new information [21].

In contrast, GPCR-Nexus fully grounds every claim in retrieved evidence from its indexed corpus, structured graph, or offline curated database. Empirical benchmarks show that Retrieval-Augmented Generation (RAG) architectures can reduce hallucinations by over 50 % compared to LLM-only approaches and improve factual consistency through explicit source retrieval [4,5]. Our Reviewer agent uses GPT-4o-mini deterministically (temperature 0) to fact-check each snippet before synthesis, guaranteeing reproducible, citation-rich answers. Additionally, the offline database returns references for its queries, which further enhances the accuracy of its answers. Unlike static chatbots, GPCR-Nexus can continually ingest new GPCR literature into its vector index and knowledge graph, ensuring up-to-date responses.

## Methods

### Document Ingestion and Text Splitting

All incoming GPCR studies (PDFs) are uploaded into a dedicated Azure Blob Storage container. A watch-triggered pipeline then invokes Azure Cognitive Search’s indexer skillset to extract raw text and metadata (title, authors, DOI) while avoiding reprocessing by marking each PDF with a processed timestamp. The Text Split skill runs in “paragraph” mode, detecting boundaries via line breaks, indentation, and punctuation, and divides each document into coherent segments of ~500 tokens. If a token-limit split falls within a paragraph, the split is deferred to the next boundary to preserve semantic integrity. To maintain context across chunks, successive segments overlap by ~50 tokens. Each chunk is annotated with document ID, page number, and paragraph index before being forwarded to downstream workflows.

### Vector Indexing

Each pre-split text chunk is embedded using OpenAI’s *text-embedding-3-small* model, producing a 1536-dimensional vector. These vectors, along with chunk metadata (ID, content excerpt, entities), are stored in Azure Cognitive Search. The indexer also applies the Key Phrase Extraction skill to tag biomedical terms (e.g., GPCR names, ligands) in each chunk. For retrieval, user queries are likewise embedded and submitted in a hybrid vector+keyword search. Azure leverages the Hierarchical Navigable Small World (HNSW) algorithm for approximate nearest-neighbor search, organizing vectors into layered graphs that yield high-recall results with low latency. By default, the top five most similar chunks are returned, each with its text, key entities, and citation.

### Knowledge Graph Construction

Simultaneously, the PDFs being processed are split into paragraph-level chunks and are fed into the knowledge-graph pipeline. A LangChain workflow uses custom prompts to drive GPT-4o-mini (temperature = 0) in extracting structured entities (e.g., “GPCR receptor,” “ligand,” “pathway,” “publication”) and relations (“binds,” “activates,” “expressed_in,” “supported_by”) from each chunk. The model outputs a JSON list of nodes and edges, which is ingested into an Azure Cosmos DB graph via the Gremlin API. Insertion logic checks for existing vertex or edge IDs to avoid duplication. Edge properties store citation contexts and snippet references. This continually growing graph makes explicit the semantic relationships among GPCRs, ligands, pathways, and literature.

### Integrated Ingestion Pipeline: From Raw PDFs to Vector and Graph Representations

Figure 1 provides an integrated overview of the ingestion pipeline that underlies the three subsections above (Document Ingestion and Text Splitting, Vector Indexing, and Knowledge Graph Construction). In practice, every new PDF uploaded to the system passes through a coordinated set of steps that transform raw text into structured knowledge that can be queried. As described in Document Ingestion and Text Splitting, each study is extracted, cleaned, and divided into semantically coherent text chunks, while metadata such as titles, DOIs, and paragraph indices are preserved. These chunks then enter the Vector Indexing workflow, where embeddings are generated, biomedical entities are tagged, and the results are stored in a high-performance vector database to enable rapid semantic search. At the same time, the Knowledge Graph Construction workflow organizes the same content into structured relationships, with gpt-4o extracting entities and edges (e.g., “ligand binds receptor”) that are inserted into the Cosmos DB graph. Together, these steps ensure that both the fine-grained narrative insights and the structured relational knowledge are captured. The figure illustrates how these streams converge: once a PDF has been successfully processed, it is simultaneously available in the vector index for semantic retrieval and in the knowledge graph for relational queries, providing the dual foundation on which GPCR-Nexus answers are built.

**Figure 1:**
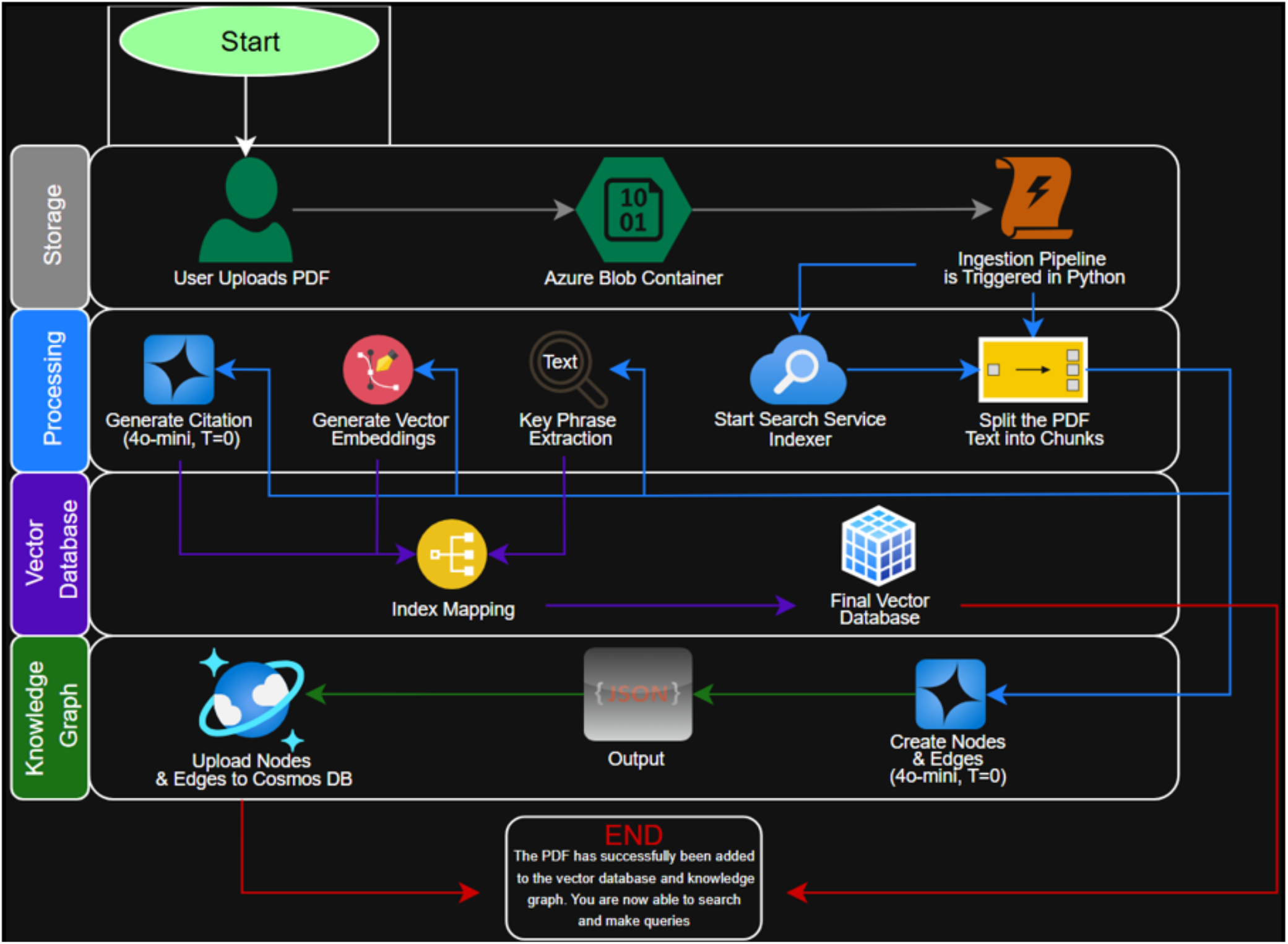
Data Ingestion Pipeline. This schematic illustrates the architectural framework designed to transform unstructured GPCR literature into a query-ready, dual-format knowledge base. Incoming PDFs are uploaded to Azure Blob Storage and processed via Azure Cognitive Search, where text is partitioned into ~500-token segments with a 50-token overlap to maintain semantic integrity. The pipeline splits into two parallel streams: (i) a Vector Indexing path, where chunks are embedded using the text-embedding-3-small model (1536 dimensions) and stored in an HNSW graph for high-recall semantic search; and (ii) a Knowledge Graph construction path, where a GPT-4o-mini workflow extracts structured entities and relations into an Azure Cosmos DB via the Gremlin API. This dual representation addresses the limitations of fragmented data by enabling both high-precision relational queries and the retrieval of narrative, mechanistic context from the literature. The pipeline ensures that every processed document is simultaneously searchable via vector-based similarity and graph-based logical connections.

### Offline Reference Database

To improve determinism and provide a more inspectable grounding layer during inference, GPCR-Nexus incorporates an offline reference database that stores structured receptor-centric profiles in SQLite. Rather than relying exclusively on downstream LLM reasoning steps to infer ligand-receptor relationships from retrieved vector database context and knowledge graph relations, the system now consults a local database containing curated receptor entries, ligand associations, synonyms, and supporting literature metadata. This architectural decision to include this makes the answer-generation process easier to audit and benchmark and reduces run-to-run variation without sacrificing query-specific context.

The offline reference database is constructed in two stages. First, an upstream data-collection pipeline retrieves human GPCR information from public structured resources, primarily IUPHAR/GtoPdb and UniProt, and assembles a reusable JSONL snapshot containing one structured receptor object per target. These objects include the canonical receptor symbol, receptor synonyms, endogenous ligand entries, additional interaction information where available, and literature-linked metadata such as PMIDs, evidence notes, and caveats. Second, the JSONL snapshot is normalized into a SQLite database with separate tables for receptors, receptor synonyms, ligands, interactions, and interaction references. This normalized structure supports fast symbol-based and synonym-aware lookup while preserving traceability to the original supporting references.

At runtime, GPCR-Nexus queries this SQLite-backed reference layer after retrieval and evidence review of the vector database and knowledge graph have been completed. The resulting structured receptor profile is then passed into the final synthesis stage together with the reviewed textual and graph-derived evidence. In this way, the offline reference layer serves as a deterministic grounding mechanism that complements the vector index and knowledge graph rather than replacing them. The vector index remains important for retrieving narrative evidence from the literature, and the knowledge graph remains useful for explicit relational context, but the SQLite layer provides a compact factual backbone for receptor-centric answering.

### Multi-Agent Pipeline

Figure 2 illustrates the agentic orchestration pipeline invoked whenever a user submits a query to GPCR-Nexus. LangChain coordinates a modular workflow involving four specialized stages. First, the Source Planner Agent identifies the receptor focus of the query and formulates targeted retrieval requests for both the vector index and the knowledge graph. These requests are executed in parallel, with Azure Cognitive Search returning semantically relevant text chunks and Cosmos DB returning subgraphs of entities and relations associated with the queried receptor and candidate ligands. The Reviewer Agent then filters, compresses, and validates the retrieved evidence, discarding irrelevant or low-confidence material and ensuring reproducibility through deterministic GPT-4o-mini settings (temperature = 0).

**Figure 2:**
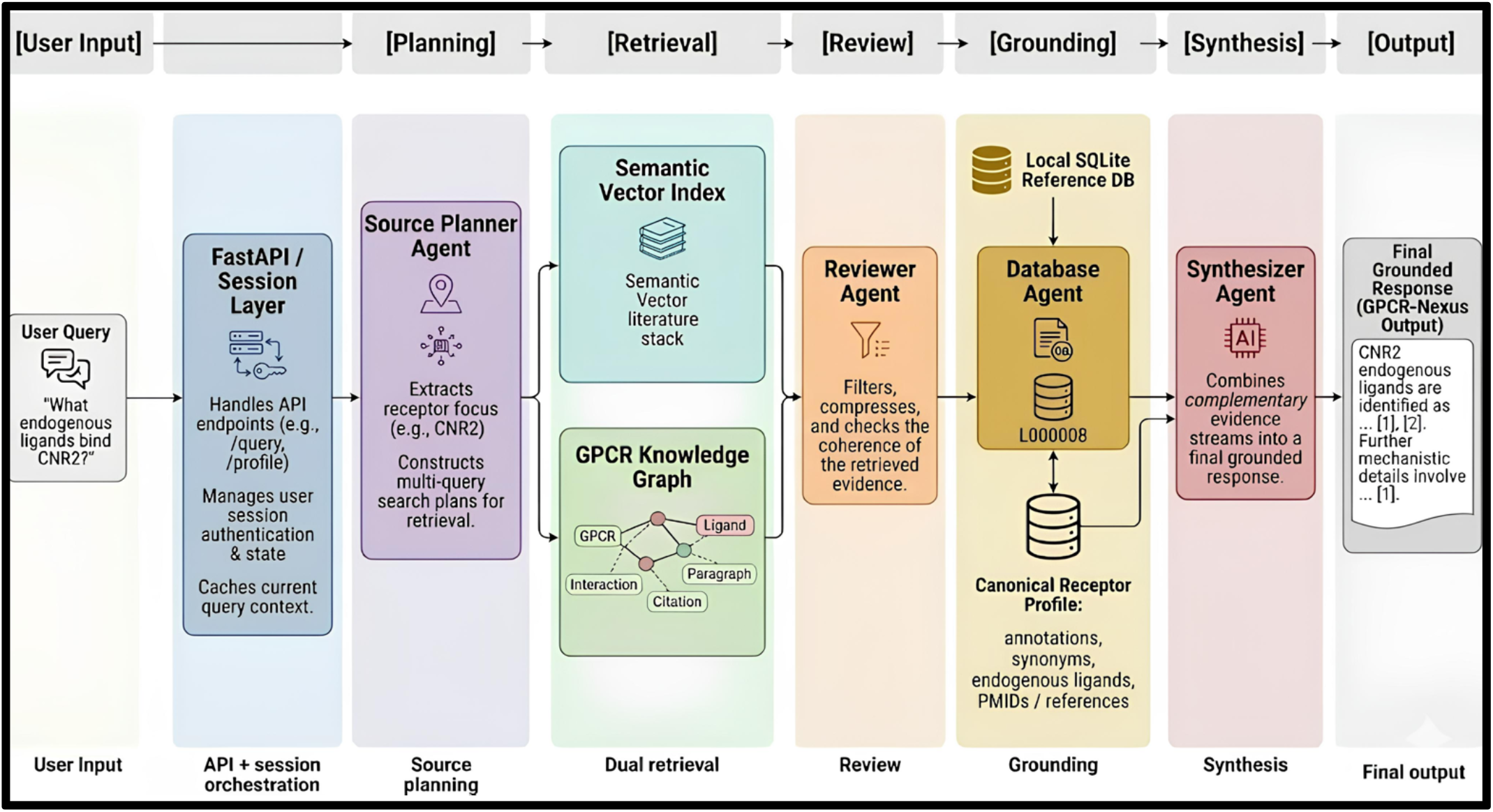
The Agentic Workflow for Knowledge Extraction and Output Synthesis. The multi-stage workflow invoked when GPCR-Nexus answers a user’s query. The Source Planner Agent extracts the receptor focus and dispatches targeted retrieval requests to both the semantic vector index and the GPCR knowledge graph. Retrieved text chunks and graph-derived relational context are then passed to the Reviewer Agent, which filters and compresses the evidence under deterministic settings to improve reproducibility and reduce irrelevant context. The reviewed evidence is subsequently augmented with a structured offline receptor profile obtained by the Database Agent from a local SQLite reference database, which contains canonical receptor annotations, synonyms, endogenous ligands, and linked reference metadata. The Synthesizer Agent then combines these complementary evidence streams into a final grounded response. This architecture allows GPCR-Nexus to integrate literature-level context, structured relational information, and deterministic local reference lookup within a unified agentic framework.

Once the Reviewer Agent has completed its steps, the Database Agent queries a local SQLite-backed receptor reference database to retrieve a structured receptor profile containing canonical naming, known endogenous ligands, synonyms, and literature-linked evidence. Finally, the Synthesizer Agent integrates the reviewed retrieval evidence and the structured offline reference output into a coherent, citation-rich response.

### Benchmark design and evaluation

To quantitatively assess endogenous-ligand retrieval performance in a manner aligned with the intended use of GPCR-Nexus, we conducted a judge-free benchmark comparing GPCR-Nexus against three general-purpose LLMs: gpt-4o, Sonnet 4.5, and Gemini 2.5. The benchmark task was intentionally narrow and standardized. Each system was asked questions of the form, “What endogenous ligands bind receptor X?” This formulation was chosen to isolate a biologically meaningful retrieval task with structured ground truth, minimize prompt variation across systems, and reduce ambiguity about the expected response type. Because the benchmark targets endogenous ligand identification rather than free-form mechanistic reasoning, the task is well suited to a judge-free evaluation framework grounded in curated receptor-ligand reference sets.

The final evaluation set comprised 100 total questions: 75 answerable receptor queries and 25 synthetic unanswerable controls. The answerable items were selected to span receptors with established endogenous ligand relationships and to include both relatively canonical and less trivial targets. The unanswerable controls consisted of deliberately fabricated receptor names designed to resemble plausible GPCR nomenclature. These synthetic controls were included to directly probe hallucination and abstention behavior under conditions where no valid receptor-ligand answer exists. We chose synthetic rather than exclusively orphan-receptor negatives because synthetic names provide a clearer negative-control condition: unlike orphan receptors, which may have incomplete or evolving biology, fabricated receptors are unambiguously unanswerable and therefore provide a stricter test of unsupported answer generation.

The comparison was intentionally asymmetric at the system level. GPT-4o, Sonnet 4.5, and Gemini 2.5 were evaluated in a closed-book setting: no browsing, no retrieval, no tools, and no external databases. They were instructed to answer using internal training knowledge only and to abstain explicitly if uncertain. By contrast, GPCR-Nexus was evaluated with its full architecture enabled, including retrieval and the local offline reference layer, because the goal of the benchmark was not to compare raw base-model memorization but to evaluate the end-to-end performance of a domain-specific GPCR question-answering system against frontier closed-book LLM baselines. This design reflects the practical question of interest: does a specialized G-protein coupled receptor agentic system outperform strong general-purpose models on structured biomedical tasks?

Gold-standard ligand-receptor relationships were assembled from curated resources, including IUPHAR/GtoPdb, UniProt, and GPCRdb, and were treated as the benchmark reference set. These sources were selected because they provide complementary strengths: IUPHAR/GtoPdb offers pharmacologically curated ligand-receptor relationships, UniProt provides structured protein-level annotations and nomenclature support, and GPCRdb offers receptor-focused coverage and cross-resource consistency. The benchmark was therefore designed to test not only nominal correctness, but also completeness of ligand-set recovery and safe behavior on unanswerable inputs.

### Manual annotation and scoring

Model outputs were manually annotated using prespecified binary indicators for full correctness, partial correctness, and hallucination (incorrectness) on (un)answerable items. This judge-free approach was used to avoid dependence on an LLM-as-judge framework and to anchor evaluation directly to a curated gold standard. Full correctness for answerable items required recovery of the *full* endogenous ligand set without introducing unsupported ligand claims. Partial correctness was operationalized as recovery of one or more correct endogenous ligands with incomplete coverage of the full ligand set, provided that unsupported ligand assertions were not introduced. Hallucination was defined as the inclusion of unsupported content inconsistent with the benchmark reference set. For unanswerable items, correct abstention, and thus no hallucination, required an explicit and appropriate acknowledgment that no valid endogenous ligand answer could be provided or exists.

Because the biological task often involves receptors with more than one accepted endogenous ligand, a strict all-or-none accuracy metric alone would understate potentially meaningful differences in answer completeness. We therefore used a graded scoring framework in addition to strict full-correct accuracy. In the primary analysis, answerable items received a score of 1 for full correctness, 0.5 for partial correctness, and 0 otherwise. Unanswerable items received a score of 1 for correct abstention and 0 otherwise. This framework was chosen to distinguish models that retrieve at least some biologically relevant signal from those that fail completely, while still preserving a strong incentive for complete ligand-set recovery. In other words, the scoring scheme recognizes partial biological utility without treating partial retrieval as equivalent to a fully correct answer.

This choice is particularly important for endogenous ligand benchmarking because many apparent “near misses” arise from incomplete ligand-set recovery rather than wholly incorrect receptor assignment. A model that names only the major ligand for a receptor with multiple endogenous ligands is qualitatively different from a model that hallucinates unsupported ligands or fails to recognize the receptor altogether. The graded score therefore provides a biologically and benchmark-relevant compromise between strict exact-match scoring and overly permissive relevance-style evaluation.

### Statistical analysis and robustness assessment

Model-level performance was summarized using the mean graded score across benchmark questions. Uncertainty in these mean estimates was quantified using nonparametric bootstrap 95% confidence intervals obtained by resampling questions with replacement. Because all models were evaluated on the same fixed set of benchmark items, model-to-model comparisons were performed using paired, question-level analyses with GPCR-Nexus prespecified as the baseline comparator. For each comparator model, we computed the mean paired difference in graded score (model − GPCR-Nexus) across matched questions and estimated bootstrap 95% confidence intervals for this paired difference. This framework was chosen because it controls for question-specific difficulty and provides a more efficient and interpretable comparison than unpaired model-level summaries alone.

To test whether observed paired differences were consistent with chance under the null hypothesis of no mean paired difference, we used two-sided paired permutation tests. Holm adjustment was then applied across the three primary pairwise comparisons to control the family-wise error rate. We selected a paired permutation framework because the outcome is discrete and benchmark sizes are modest relative to asymptotic settings. This approach avoids unnecessary parametric assumptions while remaining closely aligned with the matched-question structure of the benchmark.

Because the interpretation of partial correctness can materially affect graded metrics, robustness to the choice of partial-credit weight was assessed using a formal sensitivity analysis. Specifically, we varied the partial-credit weight continuously from 0 to 1 and recomputed graded performance and paired differences at each value. This analysis was included to determine whether the observed performance advantage of GPCR-Nexus depended narrowly on the choice of partial-credit weight or remained stable across a plausible range of scoring assumptions. Confidence intervals for the sensitivity curves were again estimated using paired nonparametric bootstrap resampling.

## Results

Across the 100-question benchmark, GPCR-Nexus achieved the highest overall graded correctness among the evaluated systems (Figure 3A). The comparator models—gpt-4o, Sonnet 4.5, and Gemini 2.5— showed lower mean graded scores, with overlapping but consistently lower point estimates than GPCR-Nexus. The distribution of discrete per-question outcomes further indicated that GPCR-Nexus produced the largest proportion of fully correct responses and comparatively fewer lower-scoring outcomes than the closed-book frontier-model baselines (Figure S1). Thus, the aggregate advantage of GPCR-Nexus did not arise from a small number of unusually high-scoring questions, but from a broader shift in the distribution toward full correctness.

**Figure 3:**
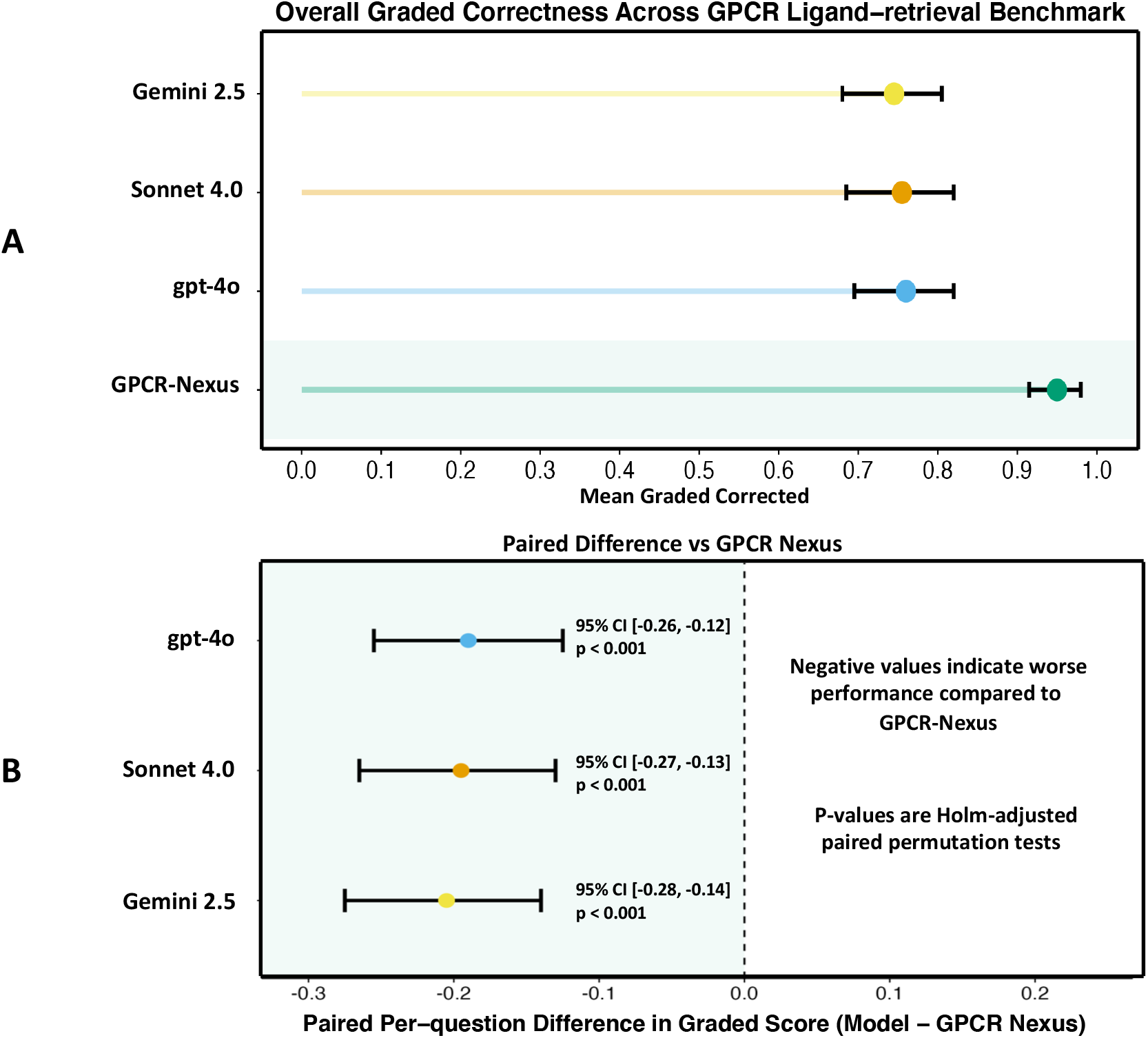
Graded Correctness and Paired Performance Differences Relative to GPCR Nexus. (A) Mean graded correctness across benchmark items for GPCR-Nexus, GPT-4o, Sonnet 4.5, and Gemini 2.5. Under the scoring rubric, fully correct answers received a score of 1, partially correct answers received a score of 0.5, and incorrect or non-creditable answers received a score of 0. Points represent model-level mean graded scores and whiskers represent 95% nonparametric bootstrap confidence intervals. GPCR-Nexus achieved the highest overall graded performance. (B) Paired per-question differences in graded score between each comparator model and GPCR-Nexus. Negative values indicate worse performance than GPCR-Nexus, and whiskers represent 95% nonparametric bootstrap confidence intervals around the mean paired difference. Holm-adjusted paired permutation p-values are annotated for each comparison. All comparator models showed negative paired mean differences relative to GPCR-Nexus, indicating that its advantage was not driven by a small number of outlier questions but was broadly reflected across the benchmark.

Paired, question-level comparisons using GPCR-Nexus as the prespecified baseline showed negative mean paired differences for all three comparator models (Figure 3B), indicating lower graded performance on the same benchmark items. The paired difference relative to GPCR-Nexus was negative for GPT-4o (95% CI approximately −0.26 to −0.12), Sonnet 4.5 (95% CI approximately −0.27 to −0.13), and Gemini 2.5 (95% CI approximately −0.28 to −0.14), with Holm-adjusted paired permutation p-values < 0.001 in all three comparisons. These paired analyses highlight the performance advantage of GPCR-Nexus is preserved under question-matched comparison rather than being driven by differences in item composition or a few outlier questions.

Sensitivity analysis further clarified the structure of these performance differences (Figure 4). As the weight assigned to partially correct answers increased, the magnitude of the paired gap between GPCR-Nexus and the comparator models attenuated but remained below zero across the full range examined. This pattern indicates that part of the GPCR-Nexus advantage derives from more frequent conversion of partially correct retrieval into fully correct ligand-set recovery. Put differently, the frontier models often retrieved some relevant ligand information, but less consistently produced the complete answer sets that was captured by GPCR-Nexus and defined by the benchmark.

**Figure 4:**
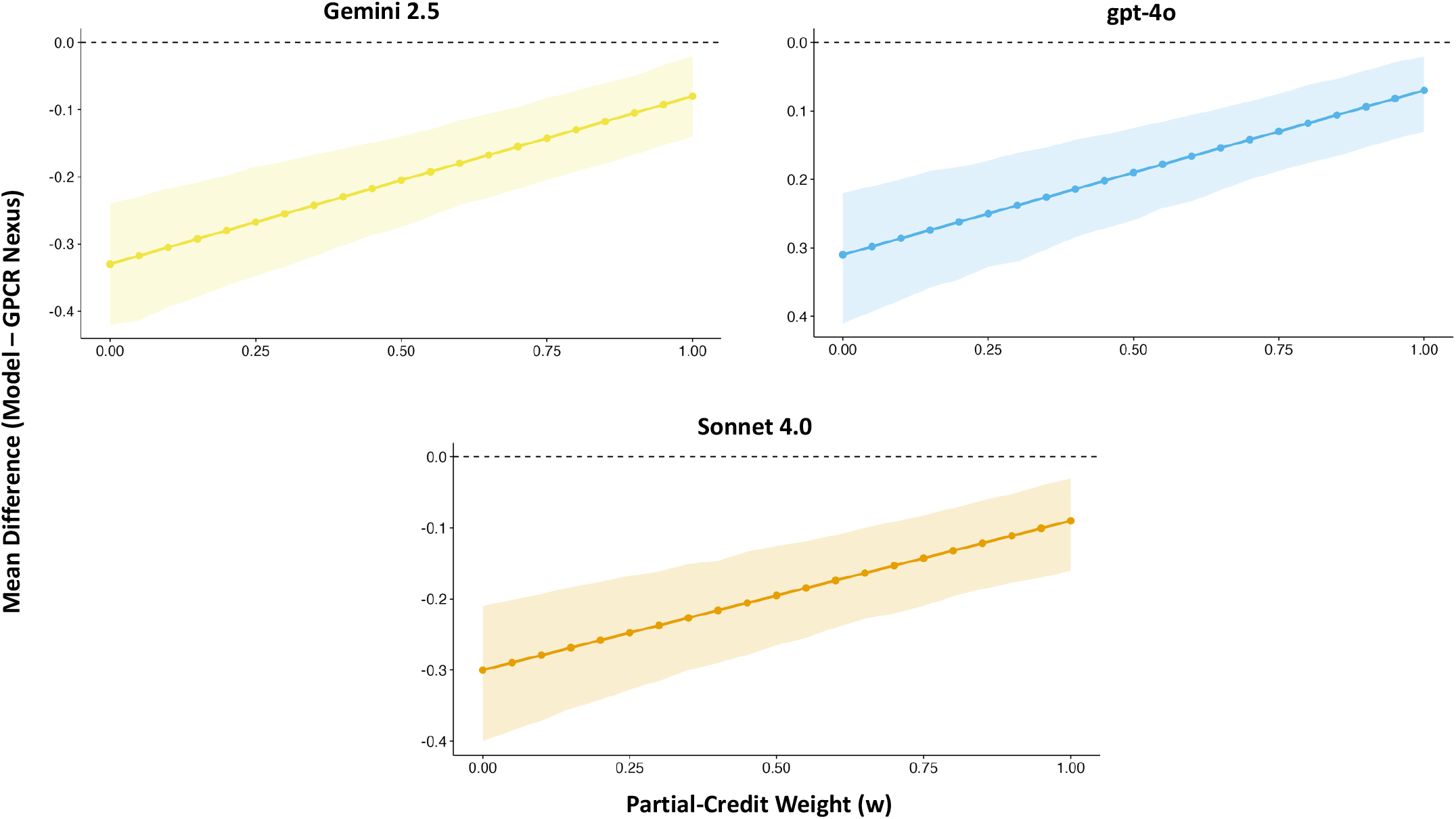
Robustness of Paired Performance Differences to Partial-credit Scoring. Sensitivity analysis assessing the stability of model performance gaps as a function of the partial-credit weight *w* ∈ {0, 0.25, 0.5, 0.75, 1.0}. The Y-axis represents the Paired Mean Difference in graded correctness relative to GPCR-Nexus; the horizontal dashed line at zero represents congruence. Shaded regions indicate 95% nonparametric bootstrap confidence intervals. For GPT-4o, Sonnet 4.5, and Gemini 2.5, the paired difference remains negative throughout the explored range, indicating that GPCR-Nexus retains an advantage regardless of how partial credit is defined. The progressive narrowing of these gaps as the partial-credit weight approaches 1 suggests that comparator models often recover some relevant ligand information but less frequently achieve the complete answer sets captured by GPCR-Nexus. Thus, GPCR-Nexus retains an advantage regardless of how partial credit is defined. The progressive narrowing of these gaps as the partial-credit weight approaches 1 suggests that comparator models often recover some relevant ligand information but less frequently achieve the complete answer sets captured by GPCR-Nexus.

Stratified analyses of answerable and unanswerable items should be interpreted with more caution than the primary paired graded-score endpoint. The benchmark was intentionally designed to include synthetic unanswerable controls to assess abstention and hallucination behavior, but these safety-related endpoints remain more sensitive to the exact construction of the negative-control set and the operational definition of non-creditable answering. Accordingly, the most robust conclusion from the present evaluation is that GPCR-Nexus outperformed the closed-book comparator models on the primary question-matched graded endogenous-ligand retrieval task, with supplementary analyses providing additional context regarding score composition and robustness to scoring assumptions.

## Discussion

These results support the hypothesis that a domain-grounded, multi-source GPCR question-answering system can outperform strong closed-book general-purpose LLMs on a specialized endogenous-ligand retrieval task when evaluated against a fixed gold standard. Importantly, this conclusion is supported not only by model-level mean graded scores, but also by paired question-level comparisons that control for benchmark-item difficulty. The paired bootstrap intervals and Holm-adjusted permutation tests indicate that the observed advantage of GPCR-Nexus is not readily attributable to a small subset of favorable questions.

The architecture also helps explain why GPCR-Nexus performed better under this task design. In the system, the answer-generation path obtains context-rich information from a vector database and knowledge graph and combines it with a deterministic offline receptor database. This makes the system more directly aligned to a receptor-ligand retrieval benchmark. However, the inclusion of the vector embeddings and knowledge graph allows for context with references to be preserved, which is just as important as discrete information answering for understanding biological importance.

The sensitivity analysis adds an important interpretive layer. The narrowing of the performance gap as the partial-credit weight approaches 1 suggests that the comparator models often retrieve a biologically relevant signal, but more frequently fail to recover complete endogenous ligand sets. This distinction matters: under a strict binary metric, incomplete yet partially informative answers can be conflated with outright failures, whereas under an overly permissive rubric, incomplete recovery can be overstated as full success. The graded analysis therefore provides a more nuanced account of model behavior and suggests that GPCR-Nexus’s advantage lies not only in avoiding incorrect answers, but also in more consistently completing the relevant ligand set.

Several limitations remain. One-hundred questions is still modest for a definitive comparative evaluation, particularly when one wishes to make fine-grained claims about individual receptor classes, ligand modalities, or abstention behavior. Second, the benchmark design is intentionally asymmetric at the system level: GPCR-Nexus was evaluated as a specialized domain agentic system, whereas the comparator models were evaluated in a generalized mode with no tool support. This asymmetry is appropriate for the system-level question being asked here, whether a domain-grounded GPCR platform improves performance over strong general-purpose baselines in G-protein coupled receptor specialized tasks. Third, scoring remains dependent on the completeness and correctness of the curated gold standard. Even high-quality resources such as IUPHAR/GtoPdb, UniProt, and GPCRdb can lag the evolving primary literature or differ in curation scope, which introduces the possibility that some apparent model errors may reflect reference-set incompleteness rather than pure hallucination. Additionally, some GPCR-Nexus answers could be marked as hallucinated if it provides a ligand for a receptor that was recently published in a paper, but hasn’t been reflected in the databases from which the gold-standard ligand-set was curated from. Fourth, the current evaluation reflects one response per system per question rather than repeated stochastic sampling, so within-model variability was not characterized.

Future work should therefore proceed in two parallel directions. On the evaluation side, larger benchmarks should expand both answerable and synthetic unanswerable items, support stratification by receptor family and ligand complexity, and incorporate multiple independent annotation passes with adjudication and inter-annotator agreement reporting. Additional analysis of ligand-set completeness, citation correctness, and calibration would further strengthen the safety and reliability claims. On the system side, the offline reference layer can be expanded to include richer provenance fields, explicit evidence hierarchies, and more systematic synchronization across curated resources. More broadly, future versions of GPCR-Nexus should distinguish clearly between benchmarking-oriented answering paths and optional hypothesis-generation modules so that factual retrieval and open-ended scientific ideation can be evaluated separately.

## Conclusion and Future Directions

In this work, we introduced GPCR-Nexus as a multi-agent, domain-grounded orchestration framework for GPCR question answering that integrates semantic literature retrieval, structured graph context, and a deterministic offline receptor database layer. The addition of the local SQLite-backed reference database strengthens the factual grounding of the system by providing an inspectable receptor-centric lookup layer that complements the provided context of the vector index and the knowledge graph. This architecture is particularly well suited to narrowly structured biomedical tasks such as endogenous ligand identification, where correctness depends not only on relevance but also on completeness and safe handling of unanswerable cases.

In a judge-free 100-question endogenous-ligand benchmark, GPCR-Nexus achieved the best overall graded performance and outperformed GPT-4o, Sonnet 4.5, and Gemini 2.5 in paired question-matched analyses. At the same time, these results should be interpreted as evidence for system-level task-specific advantage rather than as an absolute ranking of underlying base-model capability. The comparator models were evaluated in closed-book mode, whereas GPCR-Nexus was intentionally evaluated with the retrieval and reference architecture that defines it as a system. Within that framing, the present results provide encouraging evidence that specialized, retrieval-grounded biomedical systems can improve factual performance over frontier general-purpose LLMs on narrowly defined domain tasks.

Looking ahead, GPCR-Nexus provides a foundation for a broader class of domain-specific agentic systems in biology and medicine. Immediate next steps include expansion of the benchmark, richer analysis of ligand-set completeness and citation validity, and continued refinement of the offline reference layer as a first-class evidence source. Longer term, the same architecture could be extended beyond GPCRs to additional target families, including kinases, ion channels, and nuclear receptors, under a unified “Drug-Nexus” framework. Equally important will be the deliberate separation of deterministic evidence-grounded answering from more exploratory modules for hypothesis generation and experimental prioritization, allowing each capability to be optimized and evaluated on its own terms.

---

**Supplementary Figure S1:**
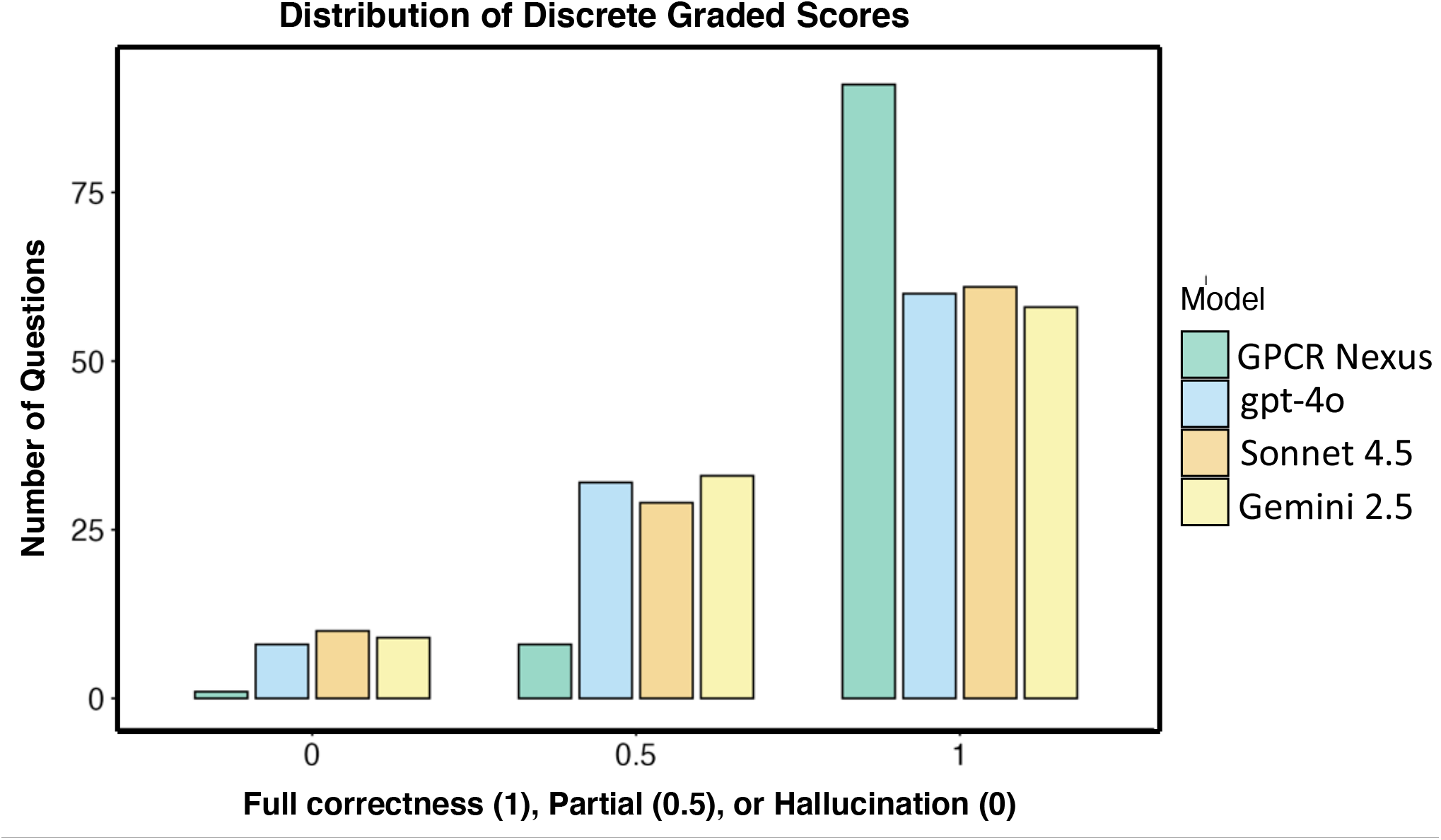
Distribution of Per-question Graded Correctness Scores Across Models Distribution of Per-question Graded Scores. Supplemental distribution of discrete outcomes across the 100-question benchmark for GPCR-Nexus, GPT-4o, Sonnet 4.5, and Gemini 2.5. Scores were assigned using the benchmark rubric in which 1 indicates a fully correct answer, 0.5 indicates a partially correct answer, and 0 indicates an incorrect or otherwise non-creditable response. GPCR-Nexus exhibits the highest number of fully correct responses and fewer low-scoring responses than the closed-book frontier-model comparators, consistent with its overall graded performance in the primary analysis. This supplemental figure provides a discrete outcome-level view of model behavior and shows that the observed advantage of GPCR-Nexus reflects a systematic redistribution toward fully correct answers across benchmark items.

## Notes

http://gpcr-nexus.org/

### Competing Interest Statement

The authors have declared no competing interest.

### Funding

- NIH Common Fund: 5R37CA272854-04
- The Center for Clinical and Translational Science and Training (CCTST)

